# Biomechanical characterization of endothelial cells exposed to shear stress using acoustic force spectroscopy

**DOI:** 10.1101/2020.09.29.319772

**Authors:** Giulia Silvani, Valentin Romanov, Charles D. Cox, Boris Martinac

**Author notes:** These authors contributed equally (Giulia Silvani, Valentin Romanov).

## Abstract

Characterizing mechanical properties of cells is important for understanding many cellular processes, such as cell movement, shape, and growth, as well as adaptation to changing environments. In this study, we explore mechanical properties of endothelial cells that form the biological barrier lining blood vessels, whose dysfunction leads to development of many cardiovascular disorders. Stiffness and contractile prestress of living endothelial cells were determined by <u>A</u>coustic <u>F</u>orce <u>S</u>pectroscopy (AFS) focusing on the displacement of functionalized microspheres located at the cell-cell periphery. The specific configuration of the acoustic microfluidic channel allowed us to develop a long-term dynamic culture protocol exposing cells to laminar flow, reaching shear stresses in the physiological range (i.e. 8 dyne cm^-2^) within 48 hours of barrier function maturation. A staircase-like sequence of increasing force steps, ranging from 186 pN to 3.5 nN, was applied in a single measurement revealing a force-dependent apparent stiffness in the kPa range. Moreover, our results show that different degrees of stiffening, defining the elastic behavior of the cell under different experimental conditions, i.e. static and dynamic, are caused by different levels of contractile prestress in the cytoskeleton, and are modulated by shear stress-mediated junction development and stabilization at cell borders. These results demonstrate that the AFS is capable of fast and high-throughput force measurements of adherent cells under conditions mimicking their native microenvironment, and thus revealing the shear stress dependence of mechanical properties of neighbouring endothelial cells.

## 1 Introduction

Cells *in vivo*, whether isolated or part of a larger collective, are constantly exposed to physical forces such as extensional, compressive, and shear stresses, all playing a critical role in regulating physiological or pathological conditions [1, 2]. Cellular mechanics and rheological properties (e.g. viscosity and stiffness) determine the ability of cells to respond to mechanical cues, and are important for many physiological processes, including growth, division and migration[3].

A remarkable example of cells under potentially high mechanical stimuli are endothelial cells (ECs) that line the inner surface of blood vessels and form the endothelial barrier. ECs form a tight tessellated monolayer which is constantly exposed to shear force generated by blood flowing along their apical surface [4, 5]. The formation and maintenance of EC contacts, which provides the functional integrity of blood vessels, requires a complex interplay of plasma membrane proteins, cytoskeletal components and associated signaling molecules [6]. ECs have evolved to form a size-selective barrier, consisting of specialized transmembrane proteins, called inter-endothelial adherens junctions, located at the cell-cell border which are ubiquitously expressed in endothelia of all vascular beds [7]. Among them, vascular endothelial cadherin (VE-cadherin), is one of the main structural and regulatory proteins controlling endothelial barrier function and permeability [8–10]. Not only do these junctions link the cells together, they also work as surface receptors generating a cascade of intracellular signaling upon external applied stimuli (e.g. shear stress), triggering dynamic interactions with cytoskeletal elements. The shear stress exerted by blood flow, typically in the range of ~1-15 dyne/cm^2^ [11–16], is known to be an important regulator of mechanotransduction in vascular physiology [17, 18], including mechanosensory processes of actin-mediated stabilization of junctions [13, 19–21], followed by morphological remodeling [6, 22–24] and tension redistribution [25] such that cells can maintain or change their shape.

According to the tensegrity model of living cells [26], the shape and stability of neighboring cells are dictated by the internal framework of contractile actomyosin filaments, better known as actin stress fibres. These active filaments are interconnected with each other and with cell-cell or cell-extracellular matrix adhesion complexes, generating a tensile prestress through a balance of complementary forces within the network [26–28]. Any variation of this structure, whether spatially or temporally, would affect the cell’s mechanical properties, e.g. cell stiffness and deformability, as well as shape stability [29–32]. In other words, the cytoskeletal prestress defines the constitutive elastic behavior of the cell and is modulated by shear stress-mediated junction development and stabilization at cell borders which in turn is strictly coordinated with actin stress fibre reorganization within the cell [32, 33]. Although the precise role of VE-Cadherin as a sensor of external stimuli remains unclear, several studies have demonstrated the importance of shear stress in *in vitro* experiments to achieve a functional monolayer with barrier properties [15, 34–39]. Besides, it is widely accepted that any alteration of the pattern of blood flow can lead to a wide range of vascular pathologies, including atherosclerosis and pulmonary arterial hypertension [40, 41]. As a result, understanding how the mechanical behavior of collective ECs may vary when exposed to fluidic shear stress is of critical importance in elucidating the cellular malfunction.

A number of micro-rheological techniques have been developed aiming at characterization of the viscoelasticity of adherent cells. These techniques include Atomic Force Microscopy (AFM), Magnetic or Optical tweezers, Micropipette Aspiration and Particle Tracking Microrheology (PTMR) [42]. Two major limitations of these techniques are measurements of reproducibility and throughput [43]. More importantly, most of the techniques require an open chamber configuration, making it difficult to mimic the physiological shear stresses necessary for EC maturation during culture. An alternative method, the recently introduced <u>A</u>coustic <u>F</u>orce <u>S</u>pectroscopy (AFS) technology, has the potential to overcome these drawbacks and greatly improve investigation of adherent cell mechanics. Originally designed for single-molecule rheology, this method uses controlled acoustic forces (in the range of pNs to nNs) over a microfluidic channel to stretch multiple molecules in parallel, individually tethered to functionalized microspheres [44, 45]. Later on, it has been used for characterization of mechanical properties of red blood cells upon different chemical treatments [46]. Recently, AFS methodology has been applied to Human Embryonic Kidney (HEK) cells capturing their inherent heterogeneity and showing the impact of temperature and pharmacological treatments on the mechanical properties at the membrane level [47].

In this study, we exploit the unique configuration of the closed AFS channel to probe the evolution of ECs mechanical properties during maturation of a confluent monolayer. By doing so, we build on the work of Nguyen et al. [48], while overcoming some reported limitations. As pointed out in their paper, viscoelastic measurements with AFS require rigorous calibration and data analysis, nevertheless a proper physiological shear stress is also needed to achieve the development of a phenotypic endothelium. First, we developed a protocol for long term, dynamic cell culture under physiological shear stress, i.e. 8 dyne/cm^2^. Second, after monolayer maturation, which was evaluated by immunofluorescence (IF) microscopy directly *in situ* at different steps, we performed creep tests by locally pulling the periphery of the cellular membrane with functionalized silica particles. The viscoelastic response of the cells was then modeled by a Power law model, to estimate the two parameters characteristic of the model, i.e. stiffness and the power-law exponent, both related to the active contractile prestress in the cytoskeleton [49]. Two different EC lines, Human Umbilical Vein Endothelial Cells (HUVECs) and Human Aortic Endothelial Cells (HAECs), were investigated to show the potential of this tool to capture heterotypic properties of endothelial barriers. Each experiment allows up to 80 measurements with forces ranging through the physiologically relevant range of pNs-nNs. Under these conditions, we demonstrated that the AFS is capable of high-throughput force measurements allowing for direct comparison between actin stress fibres reorganization, VE-Cadherin formation, and shear stress induced stiffness modulation.

## 2 Material and methods

### Experimental set-up

The AFS chip, based on a lab-on-chip device, custom fabricated by LUMICKS B.V. [44], is described in detail in SI.

### Cell culture and seeding into the AFS channel

Human Umbilical Vein Endothelial cells (HUVECs) and Human Aortic Endothelial Cells (HAECs) were purchased from Lonza (Cat. No. CC-2517, Cat. No. CC-2535). The culture medium was the endothelial basal medium-2 (EBM-2) supplemented with endothelial growth medium (EGM-2) BulletKit from Lonza (Cat. No. CC-3162). Cells were grown in tissue culture flasks and maintained in humidified atmosphere at 37°C and 5% CO_2_. The culture medium was changed every 2 days and cells were used up to the 5^th^ passage to ensure the expression of key endothelial protein components. Further details on functionalization of the AFS chip for cell seeding are given in SI.

### Immunofluorescence staining and Image analysis

Junction morphology and cytoskeleton organization was tested under different experimental conditions. VE-cadherin protein and actin stress fibres were imaged by fluorescence microscopy and ImageJ was used for scanning and analysis of the changes in morphology of both cytoskeletal proteins, as described in SI.

### Microsphere functionalization and tracking

Microspheres were functionalized with fibronectin and tracked as described in SI.

### Force steps measurement

To determine the acoustic radiation force, *F_rad_*, we performed a force-balance on acoustically driven beads [47]. The method is explained in Supplementary material together with the experimental protocol of force application. The quadratic dependence of the force with the voltage is also shown (Fig. S1).

### Data fitting

To quantify the force dependence of the creep response we used a superposition of power-law models. Details of data fitting are given in SI.

## 3 Results

The main advantage of the AFS technology is the confined fluidic circuit, shown in **Fig. 1A**. The device is also engineered and optimized to simultaneously allow for acoustic resonance and optical imaging thanks to a transparent piezoelectric element glued on top [45]. Given the small dimension of the microfluidic channels, the flow rate and the properties of the fluid, the flow is fully laminar, meaning that the flow pattern is completely predictable and does not reach turbulence (**Fig. 1Ai-ii**). Moreover, its solid structure allowed for high perfusion rates, mimicking the shear stress magnitudes exerted on ECs *in vivo*, during monolayer maturation. Here, we exploited the unique geometry of the AFS chip to perform long term culture of ECs by connecting the microfluidic channel to a peristaltic pump and maintaining the flow of growth media for 48 hours at a shear stress of 8 dyne cm^-2^. A sketch of the set-up is shown in **Fig. 1B**. The chip and the medium reservoir were placed in a dry incubator at 37 °C and 5% CO_2_ in order to avoid damaging the piezo-glass connection. During maturation, the endothelial morphology changed considerably, from an initial disorganized configuration of rounded spread cells to a compact endothelial monolayer, characterized by cell elongation in the stream direction (**Fig 1C**). Brightfield images of HAECs, after 4, 24, and 48 hours from seeding procedure are shown in **Fig 1D-F**. The elongation of cells over time in the direction of the flow is indicative for a phenotypic endothelium [50].

**Fig 1.**
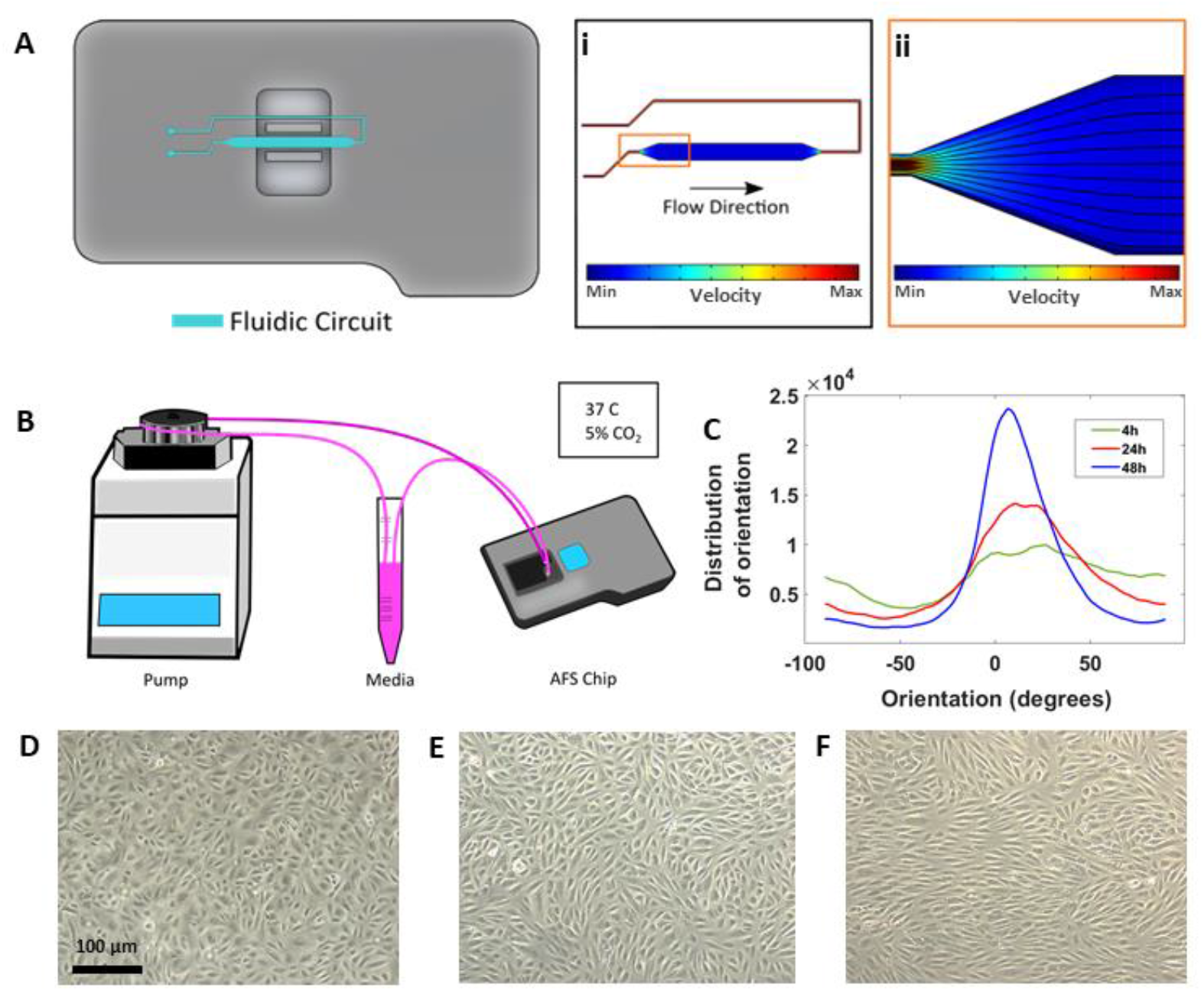
Long-term cell culture in the AFS chip. (**A**) Cartoon representation of the AFS chip (**i-ii**) Fluid dynamic simulation of fluid streamlines. (**B**) Sketch of the set up with the AFS fluidic channel connected to a peristaltic pump. (**C**) Distribution of EC orientations after 4 h (green), 24h (red) and 48h (blue) from seeding obtained with “orientationJ distribution” of ImageJ software. (**D**) Crop section of the channel imaged in transillumination mode, showing ECs 4 hours from seeding procedure and (**E,F**) cells after 24 and 48 hours, respectively, under flow culture condition at 37°C with 5% CO_2_ and no humidity. The flow was applied from left to right.

### Toward EC monolayer and junction maturation

It is generally proposed that the control of ECs adhesion, migration and barrier function, critically depends on the mutual interaction of VE-cadherin with actin filaments via receptor protein complexes [51, 52], as both structures are remodelled under physiological shear stress [22, 24]. Here, we performed IF microscopy, to assess the phenotype of these key protein components, for both EC lines, grown in the AFS device after 4 hours from seeding and at the end of the flow culture protocol, to demonstrate the significance of shear stress and timing in promoting cell barrier formation. In this section, we refer to HAECs staining and evaluation, although the same IF results were obtained with HUVECs (**Fig. S2**). When subjected to shear stresses, HAECs showed a varied response. The most striking one was the change in morphology, as they went from sub-confluence to complete confluence. Without any shear applied, i.e. CTRL, cells appeared larger and grew in a polygonal shape without directionality and with VE-cadherin proteins forming an irregular intermittent network (**Fig. 2A**, **upper left panel**) at the cell periphery. After 48 hours of flow, cell contact was achieved and VE-cadherin exhibited linear oriented junctional staining (**Fig. 2A**, **lower left panel**) suggesting a well-established, mature confluent endothelium, as expected of a tight cobblestone monolayer [50]. **Fig 2B** shows the fluorescence intensity profile for VE-Cadherin protein concentrated at the cell border for both experimental conditions. Under flow (blue curve), cells exhibited higher junctional localization of VE-Cadherin staining, as opposed to those grown under static conditions (red curve), where the junction was yet to form. Regarding rearrangement of actin stress fibres within the cellular cytoplasm, we performed fast and automated image analysis by monitoring changes in IF signal under both culture conditions. Phalloidin staining revealed a strict cortical organization of actin filaments, referred to as circumferential actin bundles [33] in cells subjected to shear stress (**Fig. 2A, lower middle panel**), whereas cells under static conditions exhibited more stress fibres across the cell body (**Fig. 2A, lower middle panel**). Therefore, at the end of the cell culture protocol, actin filaments aligned along the cell periphery sustaining and reinforcing junction proteins, as shown in the **lower right panel of Fig. 2A** by a linear, overlayed pattern of VE-Cadherin/Phalloidin. This result was also confirmed by quantification of actin stress fibres along the cells’ smaller axis, i.e. number of fluorescence peaks per μm (**Fig. 2C**). We found that, for both HUVECs and HAECs, the mid-range of fibre densities for cells under flow conditions decreased compared to control cells, suggesting substantial remodelling of actin filaments is necessary for junction stabilization (**Fig. 2D,E**).

**Fig 2.**
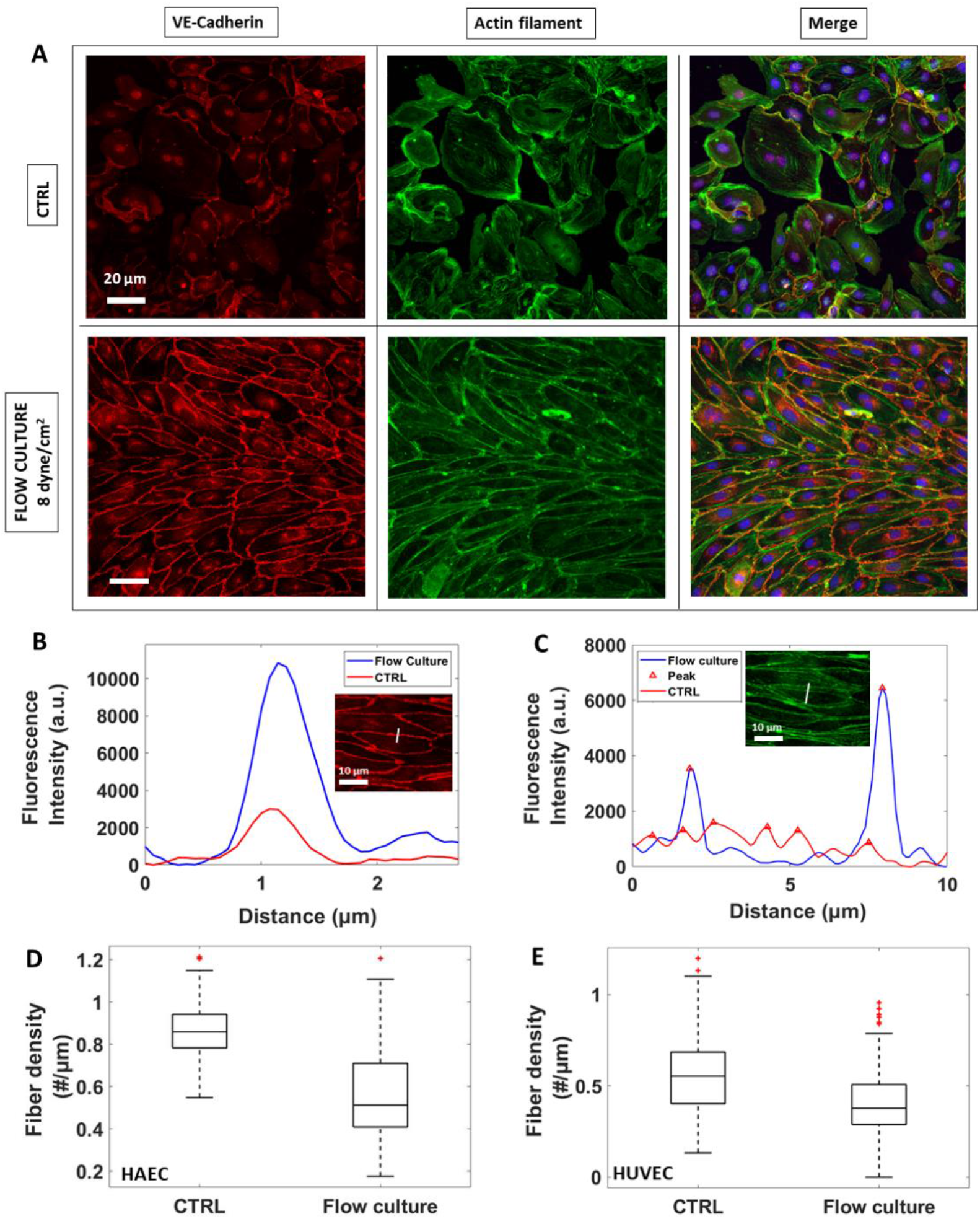
Reorganization of VE-Cadherin and actin filaments under flow culture condition. **A)** Fluorescence images of HAECs monolayer stained for junction VE-Cadherin (red, left panel) and Actin Filaments (green, middle panel) under static (CTRL) or dynamic (Flow culture) condition. **B,C)** Representative graphs showing fluorescence intensity (a.u.) profile for VE-Cadherin signal across cell-cell junctions and actin filaments, respectively. Insets: examples of the line along which fluorescence intensity profile was obtained). **D,E)** Box plots of F-actin stress fibres density for both HAECs and HUVECs.

### Manipulating endothelium monolayer with acoustic forces

In our AFS experiments, the “endothelialized” microfluidic channel was resonantly excited at 14.50 MHz, the frequency of which determines the location of the nodal plane for the acoustic standing wave. When acoustic force is applied, silica microspheres attached to the cell surface stretch the membrane of multiple adherent cells as they are pulled toward the node, capturing the cell-dependent viscoelastic creep response. The principle of an AFS experiment is shown in **Fig. 3A**. Only beads in the near proximity of cell-cell borders have been chosen for these experiments (**Fig. 3B**). In order to study the dependence of contractile prestress in the cytoskeleton with shear stress-mediated mechanical properties of neighbouring ECs, a staircase-like sequence of increasing force steps (**Fig. 3C**), ranging from 186 pN to 3.5 nN, was applied in a single measurement (i.e. differential creep test), for cells under static or dynamic conditions. Indeed, from a single-step creep experiment, no reliable conclusion about contractile prestress and shear-dependence can be made. Under the application of single stretch force, the cytoskeletal structure may not return quickly enough to its original undisturbed state in between measurements, and a large number of independent measurements is required. Besides, at higher forces, cells show a stress-stiffening response, which is consistent with adherent cell rheological behaviour in the physiologically relevant regime of large external forces and deformation [53]. We utilized the Power-Law model, 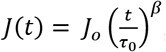, to capture the viscoelastic parameters of cells at each force step, where E_0_, the inverse of the prefactor *J_o_*, is the apparent stiffness (units of Pascal) and β, the power-law exponent, defining the solid- or liquid-like behaviour of the cell. The displacement of beads at each force step and for all cells measured, always follow a weak power-law response (**Fig. 3C,D**). In **Fig. 3E**, the cell compliance *J_o_*, decreases with increasing forces, indicating stress-stiffening behaviour.

**Fig 3.**
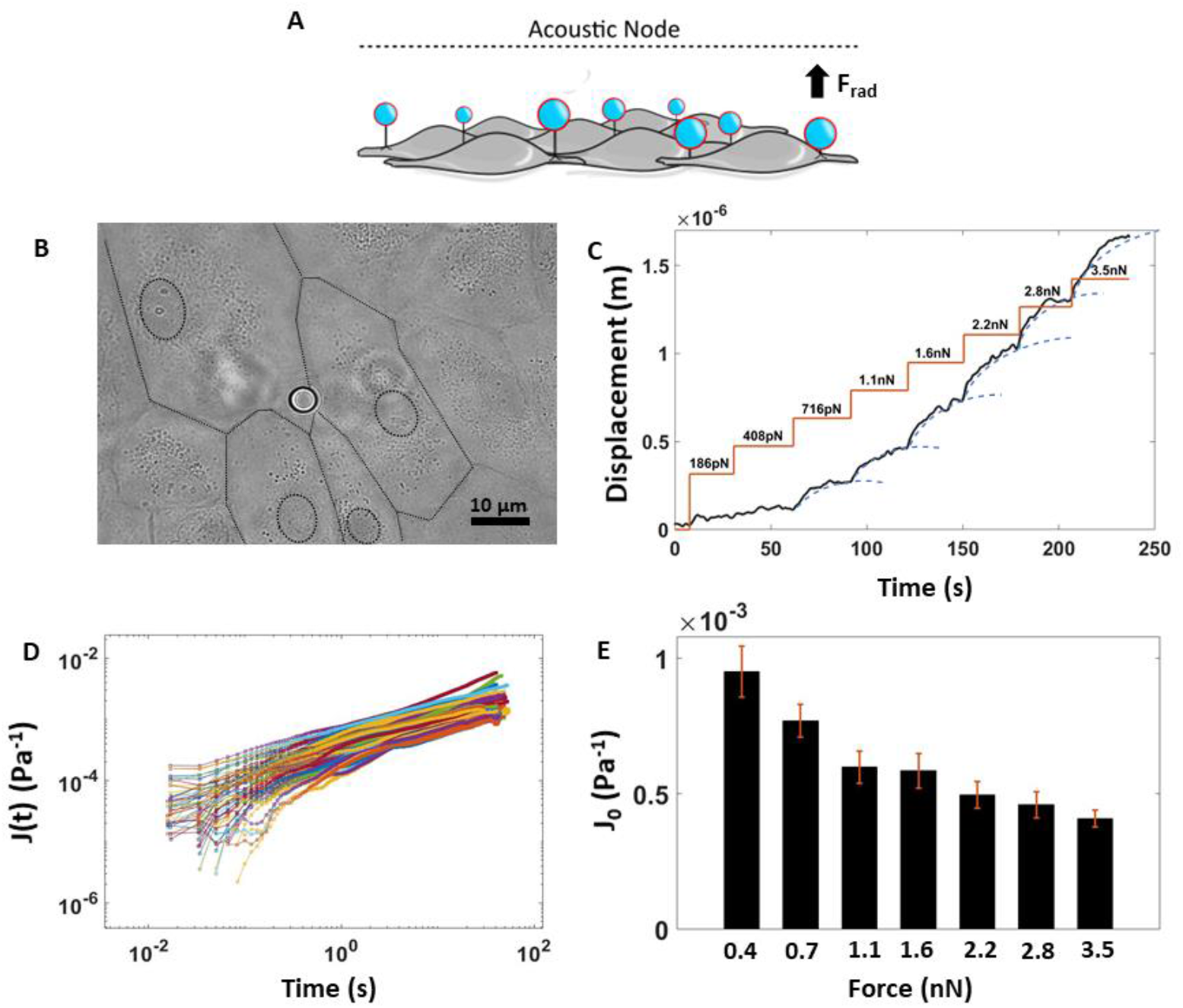
Rheology creep test for measuring nonlinear viscoelastic properties of ECs. **A)** Sketch showing the physical principle of the experiment. **B)** Bright field image of endothelium monolayer seeded in the AFS channel together with a Silica bead attached to the cellular surface. The dashed line represents the boundary of multiple cells. **C)** Creep test made of a staircase-like sequence of increasing force steps, from 180pN to 3.5nN. The bead displacement is fit to a superposition of creep responses (blue dashed line). **D)** Compliance curves at a specific force step taken from beads located all over the AFS channel and collected into a single experiment over 45 minutes (*F_rad_* = 2.8 nN, n=80, HAEC cells, EGM, 9.2 um Silica beads). **E**) Histogram showing the decrease of the factor J0 with increasing force, indicating stress stiffening, i.e. non-linear viscoelastic behaviour. Data are presented as means ± SEM.

### Local mechanical properties measured by Acoustic Force Spectroscopy

Measurements of local mechanical properties using AFS in this study indicate that different degrees of stiffening, characteristic of cells under different experimental conditions, were caused by different levels of contractile prestress in the cell, revealing the shear stress-dependence of endothelial mechanical properties.

When subjected to the staircase-like sequences of forces, both HUVECs and HAECs behaved like a viscoelastic material, in the non-linear regime, where the stiffness, 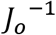, increased with increasing force (**Fig. 4A,C**). We found that 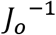 is in the order of 10^3^ Pa, consistent with other rheology measurements of adherent cells using other methods [43, 53–57]. However, variation of the power law exponent, β, was not observed (**Fig. 4B,D**). Significantly, the same stress-stiffening was found for both experimental conditions, with the only difference that the stiffness for cells under static conditions (CTRL, red curves) was higher compared to cells subjected to shear stress for 48 hours (Flow culture, blue curves). As confirmed by the stress-fibre density analysis, during monolayer maturation, actin filaments rearranged to the cell periphery by minimizing radial tension forces in the cell’s body, and thus apparently causing the cells to become softer. To further confirm this hypothesis and to find the relationship between the degree of stiffening and the prestress in the cytoskeleton, we fitted equation (S.5) to all force-stiffening curves. As shown in **Fig. 4E**, for both cell types, prestress was higher for measurements under static conditions, indicating that the tension inside the cells was not yet relaxed by the formation and maturation of protein junctions and was rather carried by contractile actomyosin filaments. These results seem to be in line with previous findings where stiffer cells exhibited higher prestress [53]. Interestingly, the prestress of HAECs was lower compared to HUVECs, while for the latter, the magnitude of the prestress under different conditions remained almost the same. This could be due to the heterotypic resistant properties of endothelium barriers from different vascular beds [58], however more experiments need to be performed in order to further confirm such hypothesis.

**Fig 4.**
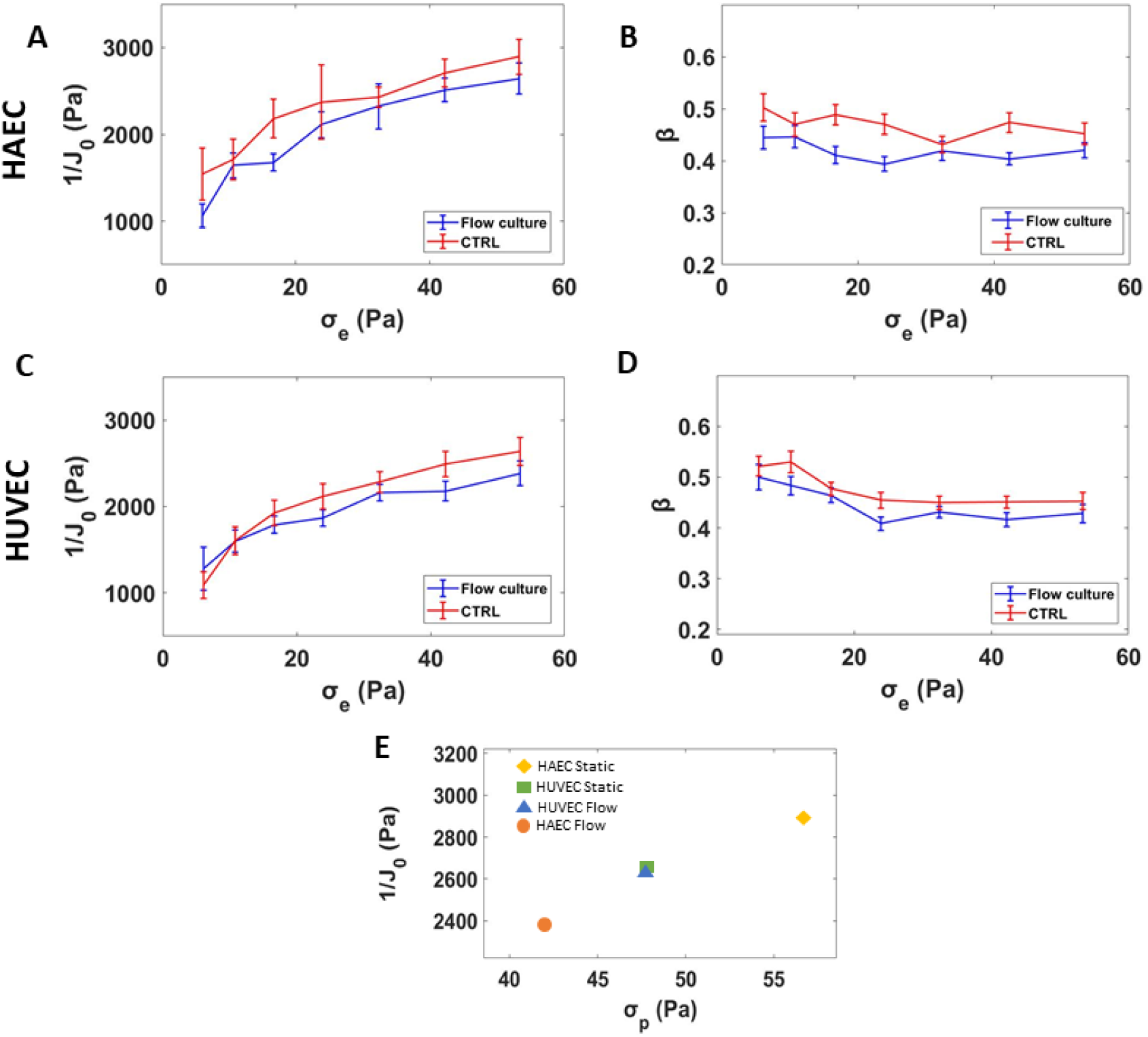
Effect of shear stress on the viscoelastic properties of ECs. **(A,C)** Stiffness, 1/J_0_, *versus* applied external stress for HAECs and HUVECs respectively, under static and dynamic condition (n=100 each). (**B,D**) The power law exponent, β, does not show any notable variation (**E**) Final stiffness *versus* prestress for all data grouped by cell type and experimental conditions. Data are presented as means ± SEM.

## 4 Discussion

The mechanisms by which ECs respond and transduce signals, such as mechanical cues, are of great importance in vascular physiology, as they are often altered during disease, driving multifunctional behavior and pathology. It is broadly accepted that the cytoskeleton of adherent cells is critical to mechanotransduction by redistributing external force across the cytoplasm e.g. shear stress from the blood. In particular, actin fibres are known to have a central role in force transmission, regulating many cellular functions, including morphological stability, adhesion, and motility [59, 60]. Although it is known that any changes in spatial distribution of actin fibres are shear dependent and occur in correlation with cadherin junction localization at cell borders, there is little quantitative knowledge about how stress fibre density and organization modulate cellular stiffness. Given that, it is important to characterize the mechanical properties of ECs under relevant physiological conditions in order to understand resulting cellular malfunction.

Much of the pioneering work characterizing adherent cells are performed using methods lacking in physiological reproduction of the cell’s native microenvironment [42]. Experimental limitations, such as an open chamber configuration, makes it difficult to assess ECs monolayer maturation under physiological shear stress conditions during cell culture. Moreover, the thickness of the cell may alter indentation measurements as it will determine the contribution from the substrate to the overall stiffness [61, 62]. Refer to [42] for a more comprehensive review on factors that may influence rheological measurements. In this study, we utilized a new technique, called Acoustic Force Spectroscopy, which operates in similar fashion to Optical Tweezers but overcomes the low throughput. Moreover, AFS technology exploits acoustic mechanical waves to manipulate a particle position, which requires simpler experimental equipment and sample preparation compared to laser set-up and usage.

After establishing a mature endothelial monolayer, assessed by IF microscopy at different growing steps, several beads at once have been pushed toward the acoustic node, stretching different cellular membranes at the surface receptors/cytoskeleton linkage site in a high-throughput fashion. By fitting a power law model to the bead’s displacement using a custom-written MATLAB program, we could determine the stiffness of ECs upon application of a range of forces and under different experimental conditions, revealing the shear stress dependence of mechanical properties of neighbouring ECs. We showed that shear stress-driven actin filament reorganization in the cytoplasm is essential for junction formation and stabilization, confirming the significance of the interplay between actin filaments and VE-Cadherin junction during barrier function maturation. Importantly, the shear stress experienced by ECs in our experiments was within the range of shear forces encountered by the cells in their native environment. This could help us to demonstrate how shear stress-driven morphology remodeling modulates cell stiffness while determining contractile prestress in the cellular body, going further with the fundamental viscoelastic measurements than Nguyen et al. [48]. Furthermore, we have extended the potential of the AFS by performing IF microscopy *in situ*, proving the applicability of AFS for further studies using live imaging. In summary, our results demonstrate the potential of AFS as a novel tool for shedding light on mechanobiology of endothelial cells, meeting several requirements during the cell culture and allowing for fast and reproducible high-throughput measurements of the viscoelastic properties at the single cell level.

## 5 Author Contributions

G.S. designed, ran, processed, and analysed the experiment and wrote the manuscript. V.R. designed, ran, processed, and analysed the experiment and edited the manuscript. C.C. helped with formulating experiments and editing of manuscript. B.M. formulated the experiments and editing the manuscript.

## 6 Supplementary Material

Supplementary material is included and contains methods for Experimental set up, Cell culture and seeding into the AFS channel, Immunofluorescence staining and Image analysis, Microsphere functionalization and tracking, Force steps measurement, Acoustic Force calibration and Data fitting.

