## Supplementary information for "Biomechanical characterization of endothelial cells exposed to shear stress using acoustic force spectroscopy"

[#]These authors contributed equally

### Experimental set up

The AFS device consists of two glass layers hosting a microfluidic channel whose thickness, width, and surface area is 100 µm, 2 mm and 54 $mm^2$, respectively. A transparent piezoelectric element glued on top allows the imaging of the sample in transillumination mode [1]. The piezoelectric element was driven with a function generator in combination with an RF-amplifier to resonantly excite a planar acoustic standing wave over the chamber (refer to *LUMICKS: AFS module* for more specification). The resonant acoustic wave was exploited to exert forces over a range of pNs-nNs on hundreds of silica microspheres (diameter, 9.2 µm, Cospheric, Cat. No. SS05003) in parallel and with sub-millisecond response time. Data were acquired using the LUMICKS AFS technology including a LabVIEW interface dedicated for microsphere tracking with an integrated temperature controller (AFS-TC, LUMICKS).

### Cell culture and seeding into the AFS channel

Here the procedure to functionalize the microfluidic chip, adapted from [2], is described in detail. Prior to cell seeding, the AFS chip was functionalized with fibronectin (100 µg/mL in EGM, Sigma-Aldrich, Cat. No F1141) using Tygon tubing (John Morris Scientific, Cat. NO ND-100-80), followed by incubation for 45 minutes at room temperature (RT). Culture flasks, with 80–90% cell confluence, were washed with Dulbecco Phosphate Buffered Saline (PBS) (Sigma Aldrich, Missouri, USA), detached using TrypLE™ solution (Gibco, Cat. No 12604-013) for 2 minutes at 37 °C in 5% $CO_2$ and blocked with growth media. Cell suspension was harvested and centrifuged at 1400 rpm for 7 min and the supernatant was discarded. Cells were resuspended in EGM at an average concentration of $10^8$ cells $ml^{-1}$ and transferred into a 1.5 ml Eppendorf tube. Using Tygon tubing, cells were then pulled through the channel up to the desired confluence of 60-70% and incubated at 37 °C + 5% $CO_2$ for 4 hours, to let the cells attach to the bottom of the AFS channels under static conditions. After incubation, the endothelialized channel was ready either for acoustic measurements or to be connected to a peristaltic pump for growth media perfusion. Initially set to 66 µlmin$^{-1}$, the flow rate was then increased to 120 µlmin$^{-1}$ (corresponding to a shear stress of 8 dyne cm$^{-2}$) until completed 48 hours of culture to achieve full junction maturation. All perfusion experiments were performed within a dry incubator to avoid damaging the AFS chip

**Immunofluorescence staining and Image analysis**

To investigate junction morphology and cytoskeleton organization under different experimental conditions,, i.e., after 4 hours (static) or 48 hours (flow culture), fluorescence imaging of VE-cadherin protein and actin stress fibre were performed. Reagents were gently pulled into the AFS channel using tygon tubing and incubated under static conditions. Cells were first washed with 1xPBS, fixed in 4% PFA for 15 min at room temperature, permeabilized for 2 min with Triton X-100 (0.5%, Sigma-Aldrich) and finally blocked with 1% BSA (Gibco, Invitrogen Corporation, Cat. No 30063-572). To monitor cell–cell contact, ECs were stained with VE-cadherin goat polyclonal primary antibody (200 mg/ml in 1% BSA) and incubated overnight. Cells were then incubated in the dark for 1 hour at RT with a cocktail solution of AlexaFluor555 conjugate-Goat anti-Mouse IgG (H+L) Secondary Antibody (2 mg/ml, Invitrogen, Cat. No A21424) and Phalloidin (1:200, Sigma Aldrich, Cat. No P5282). Nuclei were stained with DAPI. Fluorescence images of the entire device were captured with an inverted microscope Nikon Eclipse Ti2-E (specification) using Andor Zyla sCMOS camera (specification) in combination with a 20X objective (specification).

Changes in VE-Cadherin and actin stress fibres organization were evaluated by performing line scans using ImageJ and analyzing the resulting fluorescence profile. For junction maturation at the cell-cell border, lines were drawn across the contact of two cells, while for actin rearrangement, lines were drawn in individual cells, along the smaller axis, perpendicular to stress fibres. This procedure was repeated for 100 cells randomly chosen along the microfluidic channel. After correction for background, the resulting fluorescence intensity profiles were analyzed for the number of peaks above a proper baseline and at a defined distance from neighbors. In this way, two neighboring top values were considered as two separate peaks only when the distance between them was equal or higher than 0.3 µm. Dividing the number of peaks by the length of the scan line for all cells resulted in density of the actin stress fibres (#/µm). A mean density could thus be obtained.

**Microsphere functionalization and tracking**

Microspheres were functionalized with fibronectin (10 µg/mL in EGM) and incubated at RT on a rotating table for at least 1h. After incubation, the beads were injected into the endothelialized AFS channel at a cell to bead ratio of 1:2. The chip was placed on the AFS microscope stage for at least 5 minutes before being exposed to the acoustic force in order to allow for microsphere attachment to the cellular surface. At this stage of the experiment, the temperature controller was set to the physiological temperature of 37 °C. Images were acquired with a bright-field inverted microscope equipped with a 1.3 MP camera recording at 60 Hz (UI-324CP, IDS) in combination with an air 20x 0.75 NA objective (Nikon, CFI Plan APO, VC 20x, (MRD70200)). The bead z-position was determined using a predefined look-up-table (LUT) [3], a library of radial profiles as a function of z position with 100nm steps, created from a series of microsphere images prior to the application of the acoustic force.

**Force steps measurement**

To measure the non-linear creep response of cells, a staircase-like pattern of increasing constant force (ranging from 186 pN to 3.3 nN) was applied for 10 seconds at each step, for a total of 80 seconds. The displacement at each step was fitted by a creep-compliance model (see section below). All measurements were performed within 45 minutes at 37 °C with a maximum peak-to-peak driving voltage of 45 $V_{pp}$ at 14.50 MHz frequency.

**Acoustic Force calibration**

To determine the acoustic radiation force, $F_{rad}$, we performed a force-balance on acoustically driven beads. Briefly, when acoustic force is applied, beads are pushed toward the node of an acoustic standing wave. In such a scenario, the forces acting on a suspended bead in solution are the gravity force ($F_g$), the buoyancy force ($F_b$), the Stokes drag force ($F_D$) and the acoustic radiation force ($F_{rad}$). Assuming a constant velocity, all the forces at play cancel out:

$$F_g - F_b + F_D - F_{rad} = 0 \tag{S.1}$$

$$\frac{4}{3}\pi r^2 \rho_b g - \frac{4}{3}\pi r^2 \rho_m g + F_D = F_{rad} \tag{S.2}$$

where $g$ is gravity, $r$ is the bead radius, $\rho_b$ is the silica beads density and $\rho_m$ is the density of the cell media. While gravity and buoyancy forces are constant, the drag force $F_D = v_{bead}\gamma_{Brenner}$, acting on a bead moving normal to the surface, depends on bead velocity and was corrected for hydrodynamic surface effects using Brenner's factor [4]:

$$\gamma_{Brenner} = \frac{6\pi\eta r}{1-\frac{9}{8}\left(\frac{r}{h}\right)+\frac{1}{2}\left(\frac{r}{h}\right)^3-\frac{57}{100}\left(\frac{r}{h}\right)^4+\frac{1}{15}\left(\frac{r}{h}\right)^5+\frac{7}{200}\left(\frac{r}{h}\right)^{11}} \tag{S.3}$$

where η is the media viscosity and $h$ the height of the bead center to the surface [5].

The velocity was then calculated by recording the trajectory of beads moving toward the node and taking the derivative of the z-position over time. The bulk Stokes Drag coefficient, $\gamma_0 = 6\pi\eta r$ , was first inferred using Einstein relation and exploited to find the experimental medium viscosity, $\eta$, as described in [6], which was inserted in Equation (S.3) and used to calculate the effective drag coefficient. Finally, the radiation force, $F_{rad}$, could be inferred from calculating the other forces. The procedure was repeated on multiple beads (n = 17) in EGM ($\eta$ = 0.94E-3 Pa s), with increasingly applied voltage from 5 to 10 Vpp. The extrapolated forces were plotted against the applied voltage to demonstrate the quadratic dependence of the force with the amplitude (Fig.S.1).

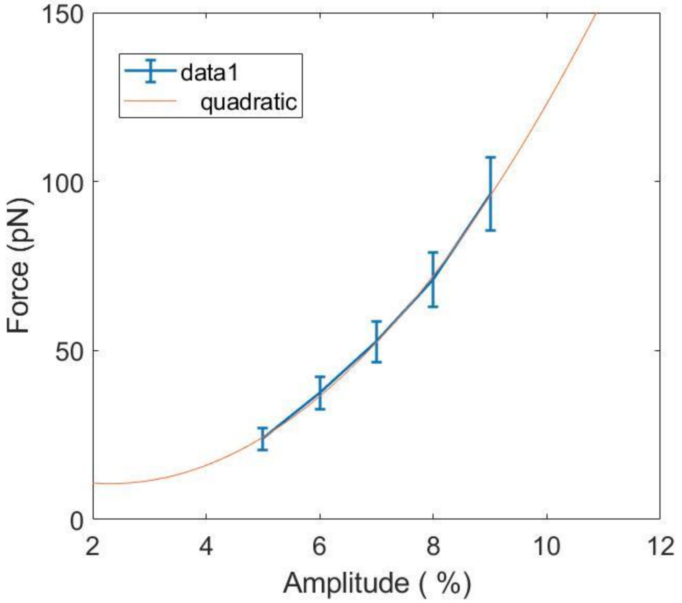


**Fig. S1.** Force calibration profile as a function of applied voltage ($V_{pp}$) for the device used in these experiments (14.51 MHz, n=17, mean±SEM) and tested with 9.2 µm Silica particles in EGM (ν = 0.94E-3 Pa s).

**Data fitting**

For each force step, we estimated the strain $\varepsilon(t)$ as the bead displacement $d(t)$ divided by the bead radius $r$ and the stress $\sigma$ as the applied force, $f$, divided by the bead cross-sectional area, $r^2\pi$. The creep compliance $J(t)$ could be determined using the expression $\frac{d(t)}{f}$ multiplied by a constant geometric factor $r\pi$ and fitted to the equation:

$$J(t) = J_o\left(\frac{t}{\tau_0}\right)^{\beta} \tag{S.4}$$

with time $\tau_0$ defined as 1. $J_o$ is the softness and $\beta$ is the power-law exponent, defining the solid- or liquid-like behavior of the cell. The value of $\beta$, falls in between 0.1 and 0.8 for most cells. The apparent elastic modulus, $E_0$, the inverse of the creep prefactor $J_o$, represents the stiffness and has a unit of Pa. The non-linear viscoelasticity (stress-stiffening behavior) was then revealed by plotting the elastic modulus, $J_o(\sigma)^{-1}$, *versus* applied stresses. With the assumption that different degrees of stiffening are caused by different levels of prestressing the cell [7], we considered the total mechanical tension of a cell's cytoskeleton to be given by the sum of active internal prestresses, $\sigma_p$, and a passive external stress, $\sigma_e$, imposed by the acoustic force. We fitted the stress-stiffness curve to the linear relationship between the stress-dependent differential stiffness $E'$, and the cytoskeleton tension, which was previously reported as a universal property of cells [8-11]:

$$E'(\sigma) = E_0' + a(\sigma_p + \sigma_e) \tag{S.5}$$

Using a single value for the prefactor $a$ and leaving $E_0'$ (linear stiffness at the force-free state) and $\sigma_p$ as free parameters, we could evaluate the contractile pre-stress for cells under different experimental conditions.

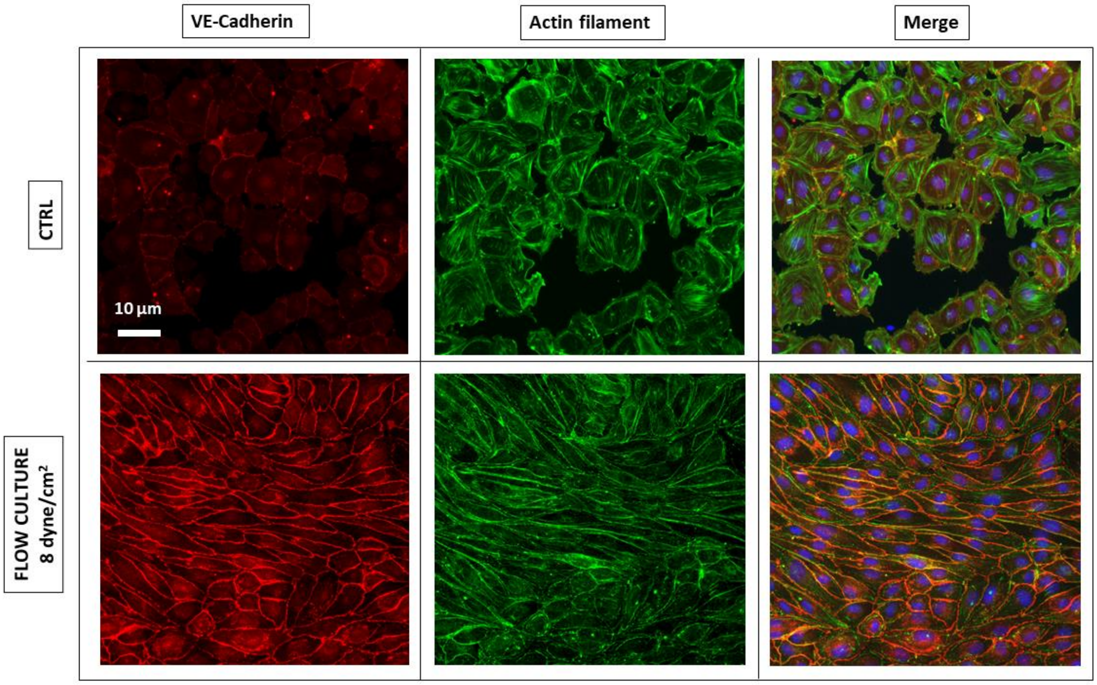


**Fig. S2.** Fluorescence images of HUVECs monolayer stained for junction VE-Cadherin (red, left panel) and Actin Filaments (green, middle panel) under static (CTRL) or dynamic (Flow culture) condition at 8 dyne/cm$^2$.


1. Kamsma, D., et al., *Tuning the music: acoustic force spectroscopy (AFS) 2.0.* Methods, 2016. **105**: p. 26-33.
2. Silvani, G., et al., *Reversible Cavitation-Induced Junctional Opening in an Artificial Endothelial Layer.* Small, 2019. **15**(51): p. 1905375.
3. Gosse, C. and V.J.B.j. Croquette, *Magnetic tweezers: micromanipulation and force measurement at the molecular level.* 2002. **82**(6): p. 3314-3329.
4. Brenner, H.J.C.e.s., *The slow motion of a sphere through a viscous fluid towards a plane surface.* 1961. **16**(3-4): p. 242-251.
5. Schäffer, E., S.F. Nørrelykke, and J.J.L. Howard, *Surface forces and drag coefficients of microspheres near a plane surface measured with optical tweezers.* 2007. **23**(7): p. 3654-3665.
6. Romanov, V., et al., *An acoustic platform for single-cell, high-throughput measurements of the viscoelastic properties of cells.* bioRxiv, 2020.
7. Kollmannsberger, P. and B. Fabry, *Linear and nonlinear rheology of living cells.* Annual review of materials research, 2011. **41**: p. 75-97.
8. Fung, Y.-c., *Biomechanics: mechanical properties of living tissues*. 2013: Springer Science & Business Media.

9. Wang, N., et al., *Mechanical behavior in living cells consistent with the tensegrity model.* 2001. **98**(14): p. 7765-7770.

10. Fernández, P., P.A. Pullarkat, and A.J.B.j. Ott, *A master relation defines the nonlinear viscoelasticity of single fibroblasts.* 2006. **90**(10): p. 3796-3805.

11. Kollmannsberger, P., C.T. Mierke, and B.J.S.M. Fabry, *Nonlinear viscoelasticity of adherent cells is controlled by cytoskeletal tension.* 2011. **7**(7): p. 3127-3132.
